# Discovery of novel Defense Regulated WD40-repeat proteins DRW1/2 and their roles in plant immunity

**DOI:** 10.1101/786848

**Authors:** Jimi C. Miller, Brenden Barco, Nicole K. Clay

## Abstract

Plant heterotrimeric G proteins transduce extracellular signals that activate plant immunity. Plants encode canonical and non-canonical Gα and Gγ subunits, but only a single canonical Gβ subunit is known. The existence of only one Gβ subunit limits the number of heterotrimeric G protein combinations able to transduce different signals. It remains unknown whether non-canonical Gβ subunits exist. Here, we identify two WD40-repeat genes that negatively regulate plant immunity. The proteins encoded by these two genes, *DEFENSE REGULATED WD40-REPEAT* 1 and 2 (*DRW1/2*), are structurally similar to AGB1. DRW2 localizes to the plasma membrane and interacts with the canonical Gα and Gγ subunits. Reduced levels of *DRW* in the *drw1* and *drw2* single mutants resulted in greater MAPK activation in response to flagellin treatment and conferred increased resistance to the bacterial pathogen *Pseudomonas syringae*. Furthermore, the *drw1 drw2* double-mutant also displayed increased MAPK activation upon flagellin treatment and broad-spectrum resistance against bacterial and fungal pathogen infection. The function of DRW1 and DRW2 is opposite of AGB1, which promotes immune signaling, suggesting that the function of these potential non-canonical Gβ subunits are not conserved with the canonical Gβ subunit. Our study identifies additional heterotrimeric G protein components, greatly increasing the number of heterotrimeric G protein complexes that participate in signal transduction.

## INTRODUCTION

Plants respond to and survive the many different environmental stresses they are subjected to, including pathogen infection. Plants perceive pathogens through the utilization of cell surface pattern recognition receptors (PRRs). PRRs bind to conserved pathogen-associated molecular patterns (PAMPs) such as flagellin from bacteria or chitin from fungal cell walls (Boller and Felix, 2009). The PRR FLAGELLIN-SENSITIVE 2 (FLS2) binds flagellin and initiates immune signaling, which in turn leads to the activation of defense responses such as the production of reactive oxygen species (ROS), activation of the mitogen-activated protein kinase (MAPK) cascade, transcriptional upregulation of pathogen-induced genes, and production of defense metabolites (Asai et al., 2002; Zipfel et al., 2004; Nürnberger and Lipka, 2005; Chinchilla et al., 2007; Boller and Felix, 2009). However, little is known about the signal transduction pathways between the receptor complex and the nucleus.

Immune signaling from FLS2 is transduced by a membrane-localized heterotrimeric G protein complex that activates downstream effectors. The heterotrimeric G protein complex is composed of a Gα, Gβ, and Gγ subunit (Temple and Jones, 2007; Oldham and Hamm, 2008). The inactive GDP-bound Gα subunit associates with the PRR and the obligate Gβγ heterodimer. When the PRR is activated, this causes a conformational change in the Gα to exchange GDP for GTP, causing the active GTP-bound Gα subunit to dissociate from the PRR complex and the Gβγ heterodimer (Temple and Jones, 2007). The Gβγ heterodimer becomes activated after dissociating from the GTP-bound Gα subunit and subsequently activates downstream effectors (Temple and Jones, 2007).

The Gβ and Gγ subunits form a Gβγ heterodimer through a coiled-coil interaction between the N-terminal α-helices on the Gβ and Gγ subunits (Sondek et al., 1996). This heterodimer is obligatory as the Gβ subunit will not fold properly in the absence of the Gγ subunit (Higgens and Casey, 1994). Interaction with the Gγ subunit promotes plasma membrane localization of the Gβ subunit through the C-terminal prenylation site on the Gγ subunit that attaches it the plasma membrane (Yasuda et al., 1996). However, there are reports in which the Gβ subunit is able to localize to the plasma membrane independently of the Gγ subunit (Ullah et al., 2008; Navarro-Olmos et al., 2010; Hackenberg et al., 2013).

Unlike animals, plants encode two different classes of heterotrimeric G proteins, canonical and non-canonical. The canonical heterotrimeric G proteins include the Gα subunit GUANINE NUCLEOTIDE-BINDING PROTEIN ALPHA-1 SUBUNIT (GPA1), the Gβ subunit ARABIDOPSIS GTP BINDING PROTEIN BETA 1 SUBUNIT (AGB1), and the Gγ subunits ARABIDOPSIS G PROTEIN GAMMA-SUBUNIT 1 and 2 (AGG1 and AGG2). The canonical heterotrimeric G proteins were discovered through homology to the animal heterotrimeric G proteins (Ma et al., 1990; Weiss et al., 1994; Mason & Botella, 2000, 2001). The non-canonical heterotrimeric G proteins include the Gα subunits EXTRA-LARGE GUANINE NUCLEOTIDE-BINDING PROTEIN 1/2/3 (XLG1/2/3), and the Gγ subunit ARABIDOPSIS G PROTEIN GAMMA-SUBUNIT 3 (AGG3). The non-canonical heterotrimeric G proteins were identified by searching for conserved domains between the non-canonical and canonical heterotrimeric G proteins, such as the Ras domain of the Gα subunit or the isoprenylation motif at the C-terminus of the Gγ subunit (Lee & Assmann, 1999; Assmann, 2002; Chakravorty et al., 2011). Despite these conserved domains, the non-canonical heterotrimeric G proteins share low homology (less than 20% protein sequence identity) to the plant canonical heterotrimeric G proteins and animal heterotrimeric G proteins (Lee & Assmann, 1999; Assmann, 2002; Chakravorty et al., 2011).

The Gβ subunit only contains two domains: an N-terminal α-helix and a seven tandem WD40-repeat domain that adopts an asymmetrical seven-bladed β-propeller-like structure (Lambright et al., 1996; Smith et al., 1999; Ullah et al., 2008; Adams et al., 2011; Ruiz et al., 2012). These two domains are not unique to Gβ subunits as there are many other WD40 repeat proteins that share a similar structure to the animal and plant Gβ subunits (Neer et al., 1994; Fulop et al., 1999; van Nocker and Ludwig, 2003; Stirnimann et al., 2010; Xu and Min, 2011). This poses a challenge in identifying novel non-canonical Gβ subunits in plants.

Both the canonical and non-canonical plant heterotrimeric G proteins are involved in plant development and immunity (Zhang et al., 2018; Liu et al., 2013; Trusov et al., 2006; Trusov et al., 2007; Xu et al., 2015; Zhang et al., 2008). Loss of either XLG2 or XLG3 results in increased susceptibility to the bacterial pathogen *Pseudomonas syringae* and the fungal pathogen *Fusarium oxysporum* (Maruta et al., 2015). Interestingly, only XLG2 seems to be involved in resistance toward the fungal pathogen *Alternaria brassicicola*, even though loss of both XLG2 and XLG3 causes severe susceptibility to all three pathogens (Maruta et al., 2015). XLG2 and XLG3 interact with the Gβγ heterodimer as well as the PRR FLS2 (Maruta et al., 2015; Liang et al., 2016). Specifically, XLG2 forms a heterotrimeric complex with the AGB1-AGG1/2 heterodimer that binds to inactive FLS2 at the plasma membrane (Liang et al., 2016). Upon flagellin binding, FLS2 initiates dissociation of the heterotrimeric G protein complex, causing phosphorylation of XLG2 to enhance the production of reactive oxygen species (ROS) (Liang et al., 2016). The Gβγ heterodimer is critical for proper immune signaling as loss of either the Gβ subunit AGB1 or both redundant Gγ subunits AGG1 and AGG2 results in increased susceptibility to *P. syringae* and the fungal pathogens *F. oxysporum*, *Botrytis cinerea*, and *A. brassicicola* (Liu et al., 2013; Trusov et al., 2007; Ishikawa, 2009; Llorente et al., 2005; Trusov et al., 2006). Both canonical and non-canonical heterotrimeric G proteins are crucial in immunity.

The discovery of non-canonical Gα and Gγ subunits raises the possibility of the existence of non-canonical Gβ subunits. However, plant genomes typically encode more than 200 putative WD40-containing proteins with a similar structure to the Gβ subunit (Ouyang et al., 2012; van Nocker and Ludwig, 2003), which poses a challenge in identifying potential non-canonical Gβ subunits. To overcome this challenge and identify non-canonical Gβ subunits that participate in immune signaling, we analyzed the transcriptional profiles of 168 genes that encode proteins with seven tandem WD40-repeat motifs for up- or downregulated expression upon biotic stress. We identified two WD40 repeat genes, *DEFENSE REGULATED WD40-REPEAT 1* and *2* (*DRW1/2*), that were transcriptionally regulated by different pathogen conditions in *Arabidopsis thaliana*. DRW1 and DRW2 are structurally similar to the canonical Gβ protein AGB1. DRW2 localizes to the plasma membrane and interacts with the canonical Gα and Gγ subunits. Gene knockdown of *DRW1* and *DRW2* resulted in broad-spectrum resistance to bacterial and fungal pathogens suggesting *DRW1* and *DRW2* negatively regulate plant immunity. However, the negative immune regulation of these potential non-canonical Gβ subunits is unexpected as the canonical Gβ subunit promotes immunity. This study provides the first evidence of novel non-canonical Gβ subunits which greatly increases the number of heterotrimeric G protein combinations that are able to function in wide array signal-transduction pathways in plants.

## RESULTS

### *DEFENSE REGULATED WD40-REPEAT* (*DRW*) gene family expression is modulated upon biotic stresses

Non-canonical Gα and Gγ subunits involved in plant immunity have recently been discovered (Lee and Assmann, 1999; Assmann 2002; Thung et al., 2012), but it remains unknown whether non-canonical Gβ subunits that function in immunity exist. Plant canonical Gβ subunits contain seven tandem WD40 repeats and are upregulated upon biotic stress (Winter et al., 2007; Lee et al., 2013). To identify potential non-canonical Gβ subunits, we identified 168 genes containing seven tandem WD40 repeats. Next, we analyzed their gene expression under different biotic stress conditions by searching publically available expression data in the Bio-Analytic Resource Expression Angler. The canonical Gβ, *AGB1*, was upregulated in response to bacterial and fungal infections and highly upregulated upon PAMP treatment (Figure 1). By searching the microarray data, we identified a gene family whose gene expression was up- or downregulated in response to biotic stressors, which we named *DEFENSE REGULATED WD40-REPEAT (DRW)*. This gene family consists of five genes (*AT1G55680*, *AT3G13340*, *AT5G56190*, *AT1G78070*, and *AT1G36070*), which we named *DRW1*, *DRW2*, *DRW3*, *DRW4*, and *DRW5*, respectively. Under PAMP treatment, *DRW1* expression was initially repressed upon flagellin elicitation and later upregulated after flagellin treatment. Moreover, *DRW1* was slightly repressed upon bacterial and fungal pathogen infection (Figure 1). Interestingly, *DRW2* showed little transcriptional change upon PAMP treatment, although it was repressed upon bacterial and fungal infection. Conversely, *DRW3* transcripts were upregulated upon bacterial and fungal infection and PAMP elicitation, which closely resembled the expression profile of the canonical Gβ subunit *AGB1*. *DRW4* was repressed upon PAMP treatment, infection by various bacterial pathogens, and infection by the fungal pathogen *Phytophthora infestans*, similar to the expression pattern of *DRW1* and *DRW2*. *DRW5* transcripts were upregulated in response to some bacterial pathogens and the fungal pathogens *P. infestans*, *Botrytis cinerea*, and *Golovinomyces orontii*. However, *DRW5* was down regulated upon PAMP treatment. Aside from *DRW3*, gene expression in this family was opposite from that of the canonical Gβ *AGB1* upon biotic stress, which is generally upregulated upon PAMP and pathogen infection. The identification of these *DRW* genes and their response to biotic stress suggest them to be potential non-canonical Gβ subunit candidates.

**Figure 1.**
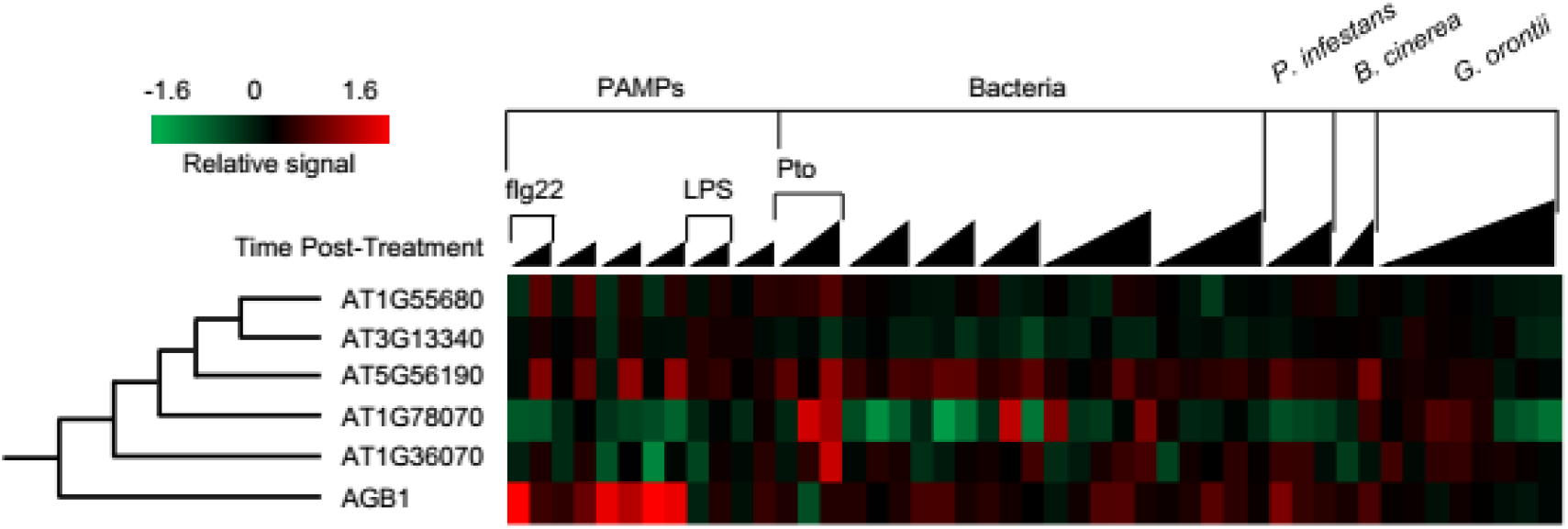
WD40 repeat protein transcripts are modulated upon biotic stresses. Gene regulation of *AT1G55680*, *AT3G13340*, *AT5G56190*, *AT1G78070*, and *AT1G36070* upon biotic stresses at various time points. Width of black triangles represent the number of time points and the height represents later time points. Data was obtained from the Bio-Analytic Resource Expression Angler in which gene expression was normalized to their respective control treatments. Then data was put on a log scale (base 2) to observe repression and upregulation of genes.

### *DRW* gene family is phylogenetically distinct from the Gβ subunit AGB1 and the WD40 RACK1 proteins

AGB1 is an established Gβ subunit (Weiss et al., 1994; Anderson and Botella, 2007). However, it remains unclear if the DRWs, a potential non-canonical Gβ protein family, is evolutionarily related to AGB1. To gauge the evolutionary relationship between the DRW protein family and AGB1, we performed a multiple sequence alignment of all 168 seven-tandem repeat WD40 proteins in the Arabidopsis genome using Clustal Omega. We generated a maximum likelihood phylogenetic tree using the Clustal Omega data. We found that the DRW protein family clustered into its own protein family, which was phylogenetically distinct from AGB1 (Figure 2A). Sequence identity analysis showed a high level of sequence identity (approximately 57%) within the DRW protein family. DRW1 and DRW2 shared the highest level of homology, with an 89% protein sequence identity (Figure 2B). Protein sequence identities between the DRW protein family, AGB1, and the human Gβ subunit GUANINE NUCLEOTIDE-BINDING PROTEIN SUBUNIT β-1 (GNB1) were below 20%. However, sequence identity between AGB1 and GNB1 was 46%. While the DRW protein family was distinct from AGB1, the DRW protein family was more related to AGB1 than other known WD40 repeat proteins such as TRANSPARENT TESTA GLABRA (TTG1) and RECEPTOR FOR ACTIVATED C KINASE 1A (RACK1A), RECEPTOR FOR ACTIVATED C KINASE 1B (RACK1B), and RECEPTOR FOR ACTIVATED C KINASE 1C (RACK1C) (Figure 2A). As these known WD40 proteins did not contain the N-terminal α-helix that facilitates the interaction with the Gγ subunit and promotes localization of the Gβ subunit to the plasma membrane, they were selected as controls for this experiment. Protein sequence identity between the DRW protein family and AtRACK1A was below 20% suggesting that the DRW protein family is unrelated to the RACK1 proteins (Figure 2B). These results provide further support that the DRW protein family may contain non-canonical Gβ subunits.

**Figure 2.**
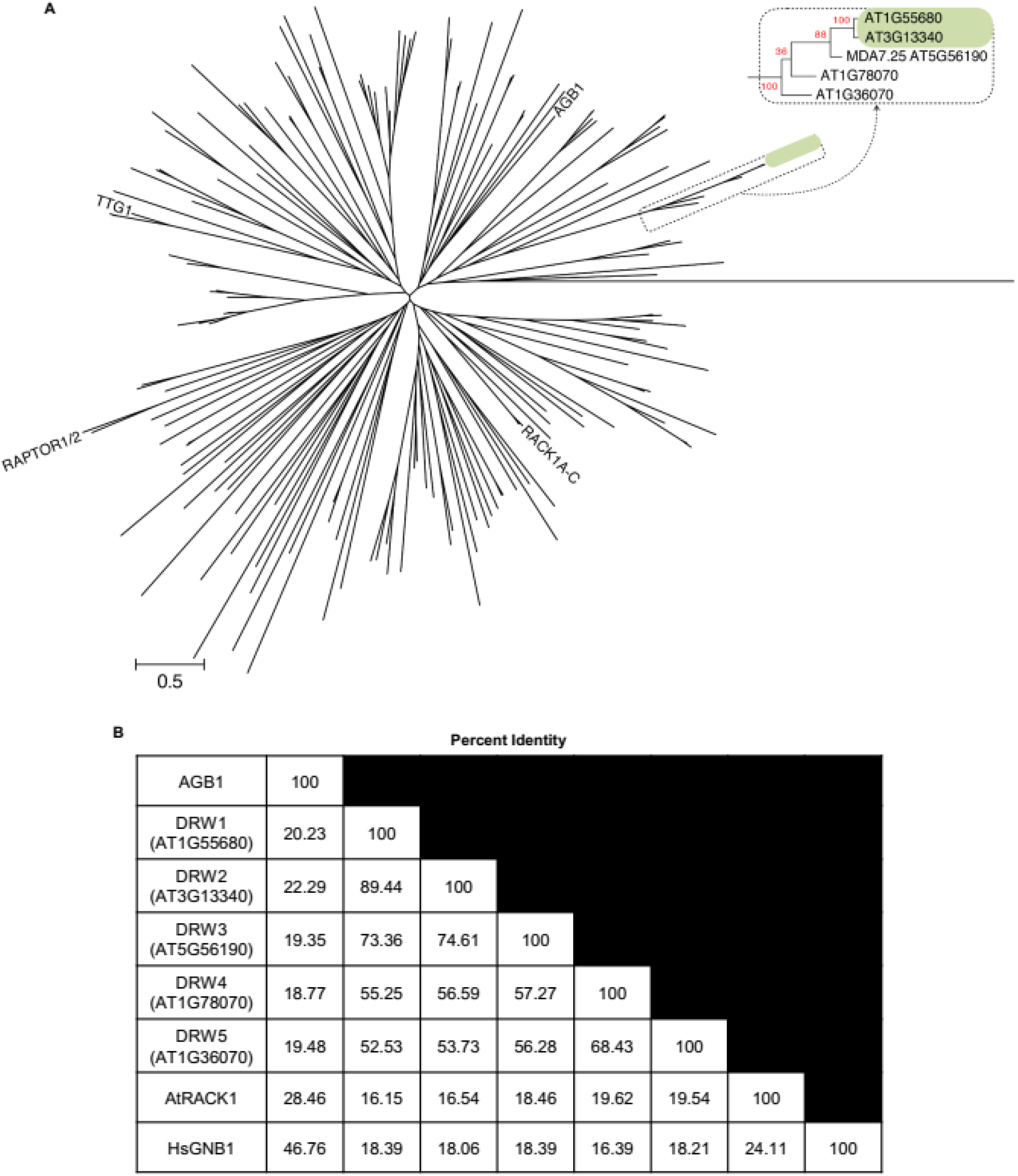
WD40 protein family is phylogenetically distinct from the Gβ subunit AGB1 and the WD40 RACK1 proteins. (A) Maximum Likelihood-based tree of 168 seven WD repeat-containing proteins based on the JTT matrix-based model in MEGA7. All amino acid positions with less than 90% site coverage were eliminated. Scale bar denotes the number of substitutions per site. (B) Percent identity matrix comparing protein sequence identity between *Arabidopsis* AGB1, DRW1 (AT1G55680), DRW2 (AT3G13340), DRW3 (AT5G56190), DRW4 (AT1G78070), DRW5 (AT1G36070), *Arabidopsis thaliana* RACK1A, and *Homo sapiens* GNB1. Protein sequences were aligned using the Clustal Omega multiple sequence alignment program.

### DRW1 and DRW2 predicted protein structures are similar to the canonical Gβ subunit

Protein structure is indicative of functional homology between two proteins. To determine if structural homology exists between AGB1 and the DRW proteins, we used the Phyre2 Protein Fold Recognition Server to predict the structure of the DRW proteins. This algorithm uses homology detection methods to predict the secondary and tertiary structures to build a 3D model of the protein of interest (Kelly and Sternberg, 2009). We chose to analyze *DRW1* (*AT1G55680)* and *DRW2* (*AT3G13340)* due to the high protein sequence identity between these two proteins, the similarity of their expression patterns in response to biotic stress, and the availability of T-DNA insertion mutants for *DRW1* and *DRW2* but not the other DRW family members. DRW1 and DRW2 were predicted to form an asymmetrical seven-bladed β-propeller with an N-terminal tail. Moreover, predicted protein structures of DRW1 and DRW2 were more similar to the canonical Gβ subunit AGB1 and the human Gβ subunit GNB1 than the Arabidopsis WD40 repeat protein RACK1A, which lacks the N-terminal tail (Figure 3A). Interestingly, DRW1 and DRW2 contained an additional 50 amino acids at their N-termini that are absent in AGB1 or GNB1, suggesting the N-terminus of DRW1 and DRW2 may have novel functions in addition to interacting with the Gγ subunit. Our protein homology results provide further support that DRW1 and DRW2 are potential non-canonical Gβ subunits.

**Figure 3.**
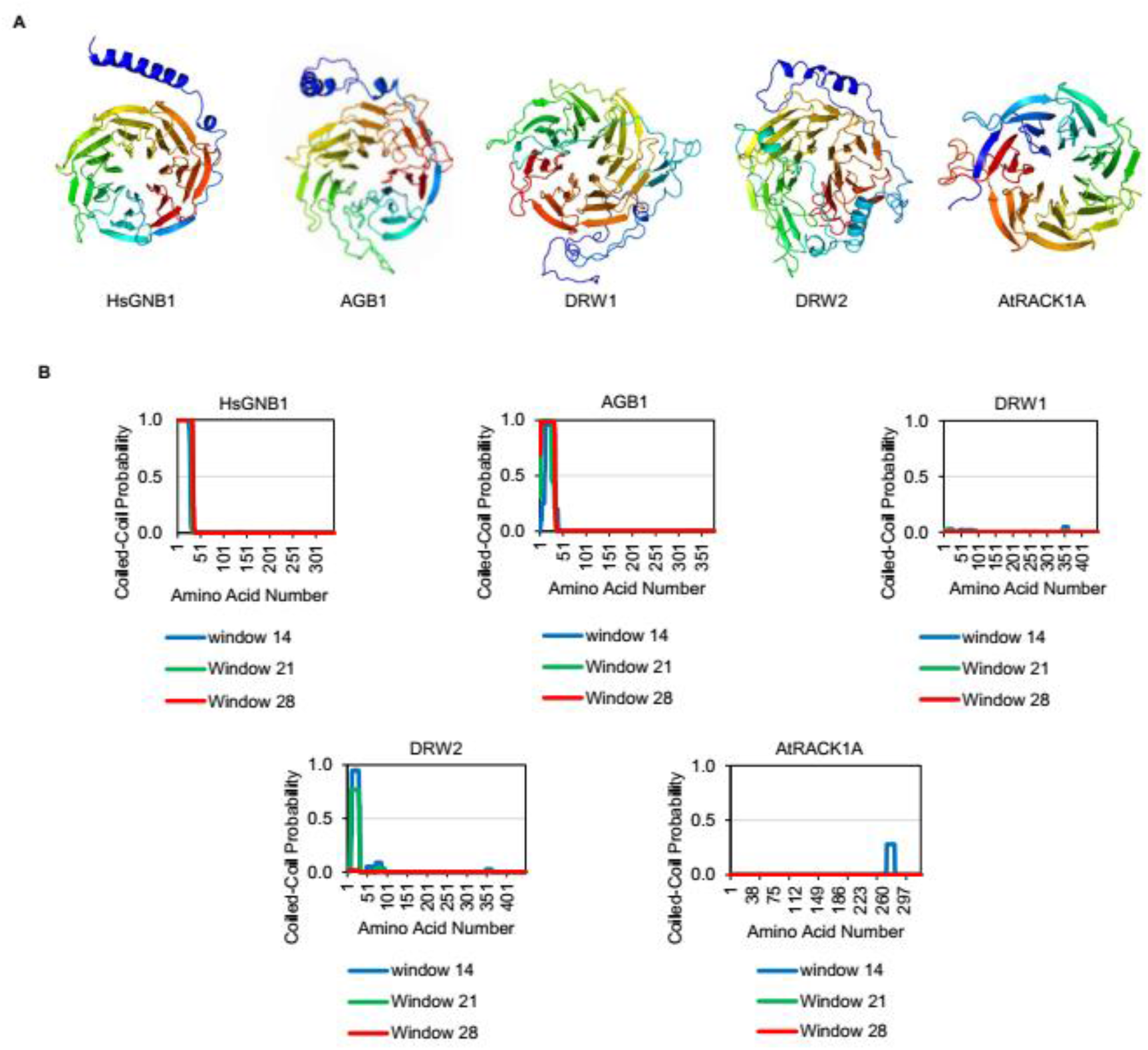
DRW1 and DRW2 proteins share a similar predicted protein structure as the canonical AGB1. (A) Protein structures were predicted using homology models based on the known structure of the *Homo sapiens* β subunit GNB1 as well as from multiple sequence templates using the Phyre 2.0 server (http://www.sbg.bio.ic.ac.uk/phyre2/html/page.cgi?id=index). Proteins from left to right: *Homo sapiens* GNB1, *Arabidopsis thaliana* AGB1, *Arabidopsis thaliana* DRW1, *Arabidopsis thaliana* DRW2, and *Arabidopsis thaliana* RACK1A. (B) Predicted coiled-coil domains of HsGNB1, AGB1, DRW1, DRW2, and AtRACK1A. Coiled-coil domains were predicted using the COILS prediction program. Windows depict three independent predictions.

DRW1 and DRW2 are predicted to have similar protein structures as AGB1. It is unknown whether the N-terminus of DRW1 and DRW2 are predicted to form coiled-coil domains to interact with the N-terminal α-helix of the Gγ subunit, as seen in the Gβγ protein crystal structure (Wall et al., 1995; Sondek et al., 1996). We used the COILS prediction program to predict if DRW1 and DRW2 were able to form coiled-coil domains at their N-termini (Lupas et al., 1991). DRW2 was predicted to form a coiled-coil domain at its N-terminus (Figure 3B). This is similar to HsGNB1 and AGB1, which are known to interact with their respective Gγ subunits through coiled-coil domains (Wall et al., 1995; Sondek et al., 1996; Obrdlik et al., 2000). Interestingly, DRW1 was not predicted to form a coiled-coil domain at its N-terminus (Figure 3B). This data suggests that DRW2 is able to form coiled-coil domains with the N-terminal α-helix of the Gγ subunit while DRW1 may not, though further functional analyses are needed.

### DRW1 and DRW2 co-localize with Heterotrimeric G protein complexes

The heterotrimeric G proteins function at and localize to the plasma membrane (Anderson and Botella, 2007), however, the subcellular localization of DRW1 and DRW2 is unknown. To determine the subcellular localization of DRW1 and DRW2, we expressed *DRW1-GFP* and *DRW2-GFP* with plasma membrane markers in tobacco leaves. DRW2 co-localized with the membrane receptor FLS2 and the Gγ subunit AGG1 at the plasma membrane (Figure 4A). Small populations of DRW1 co-localized to the plasma membrane with FLS2 whereas the majority of DRW1 did not, suggesting cytoplasmic localization (Figure 4A). In concordance with our structural studies on the DRW1 protein, DRW1 was not stably localized to the plasma membrane, reducing the likelihood that it functions as a non-canonical Gβ subunit. However, our data shows that DRW2 localizes at the plasma membrane, suggesting that it functions as a Gβ subunit.

**Figure 4.**
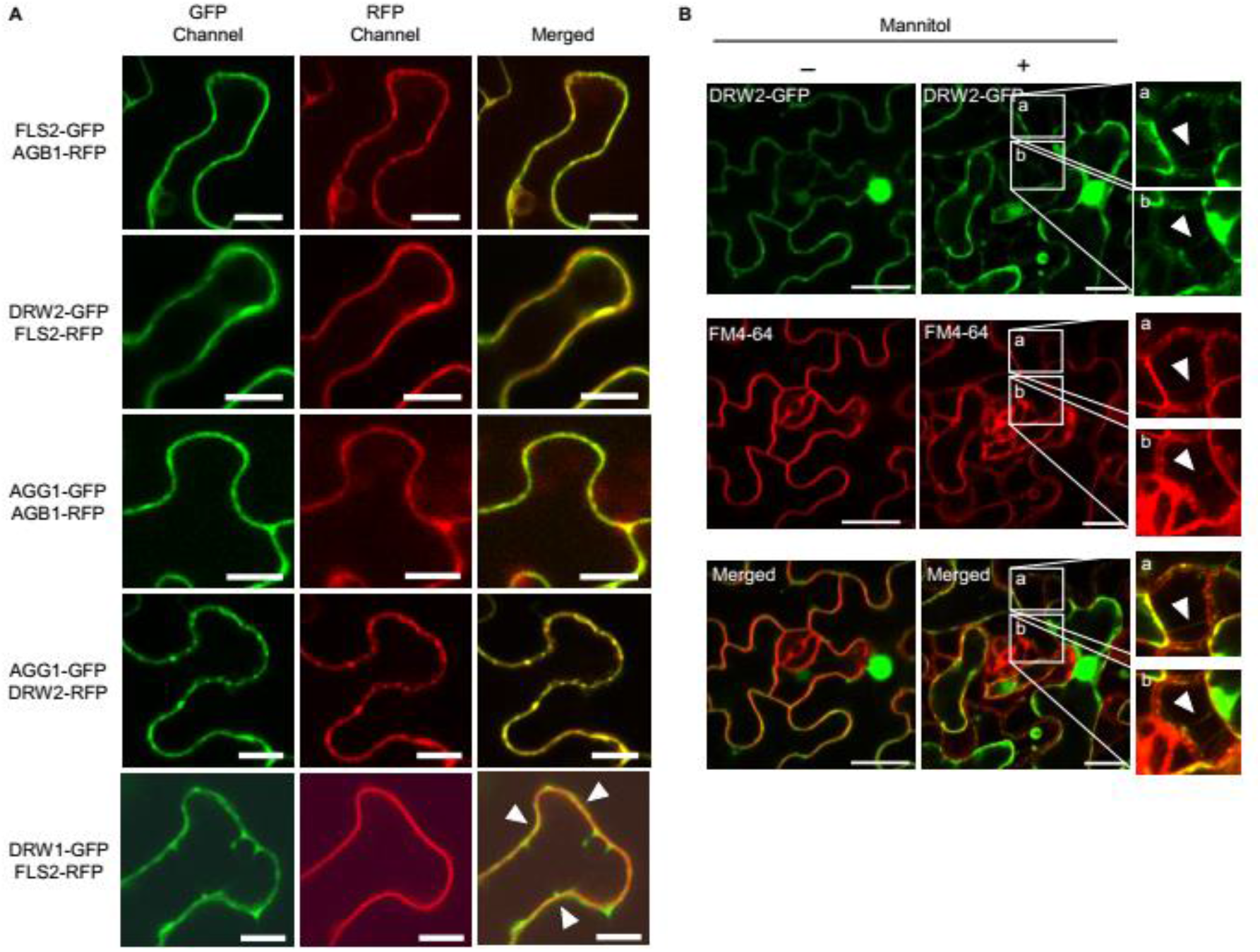
DRW2 localizes to the plasma membrane. (A) Co-localization of DRW2 with FLS2 and AGG1 at the plasma membrane. Fluorescently labeled DRW1 and DRW2 proteins were transiently expressed with 20 μM β-estradiol for 4-8 hr in *N. benthamiana* leaves. FLS2-RFP is a plasma membrane marker. White arrowheads represent populations of co-localization. White bars represent 20 μm. (B) Co-localization of DRW2 with the Hecthian strands. *DRW2-GFP* was expressed with 20 μM β-estradiol for 4-8 hr and then the plasma membrane was labeled with FM4-64. Leaf sections were imaged before and after plasmolysis with mannitol.

The plant vacuole pushes the cytoplasm against the plasma membrane, making plasma membrane localization difficult to discern from cytoplasmic localization. To validate that DRW2 localizes to the plasma membrane, we performed plasmolysis experiments to shrink the vacuole and pull the plasma membrane away from the cell wall. This leaves sections of the plasma membrane attached to the cell wall creating strands, called Hechtian strands (Buer et al., 2000). We expressed *DRW2-GFP* in tobacco leaves, labeled the plasma membrane with FM4-64, and induced plasmolysis in tobacco leaves with mannitol. DRW2 localized to Hechtian strands labeled with FM4-64, indicating DRW2 localized to the plasma membrane (Figure 4B). Taken together, these data show that small populations of DRW1 localize to the plasma membrane, although further localization experiments are required, and that DRW2 localizes to the plasma membrane thus providing additional evidence that it may be a non-canonical Gβ subunit.

### DRW2 interacts with the canonical Gγ subunits AGG1/2 and the Gα subunit GPA1

A feature of the Gβ subunit AGB1 is that it interacts with the Gγ (AGG1 and AGG2) and Gα (GPA1) subunits (Wang et al., 2008). If DRW1 or DRW2 are non-canonical Gβ subunits they would likely interact with the canonical Gα subunit GPA1 and the Gγ subunits AGG1 and AGG2. To determine the protein-protein interaction profiles of DRW1 and DRW2 with the canonical heterotrimeric G proteins, we used a split-luciferase complementation assay. The N-terminal portion of the luciferase gene was translationally fused to one putative interacting protein and the C-terminal portion of the luciferase gene was translationally fused to the other putative interacting protein. If the two proteins interacted, the two halves of the luciferase protein would reconstitute luciferase activity. When AGB1 was co-expressed with either AGG1 or AGG2, their interaction produced luminescence that was significantly higher than the control proteins expressed in Arabidopsis protoplasts. However, DRW1 co-expression with AGG1 or AGG2 showed no interaction, indicating that DRW1 did not interact with either Gγ subunit. When DRW2 was co-expressed with AGG1 or AGG2, DRW2 interacted with AGG1 and AGG2, though luminescence was not as high as the AGB1-AGG1 or AGB1-AGG2 interaction (Figure 5A). This suggests that DRW2 interacts with AGG1 and AGG2, albeit with a lower binding affinity than the AGB1-AGG1/2 interactions. AGB1 expression with GPA1 in protoplasts showed interaction, which is in accordance with a previous study showing the Gα subunit GPA1 interacts with the Gβ subunit (Wang et al., 2008). When DRW1 was co-expressed with GPA1, no luminescence was produced, suggesting DRW1 did not interact with the Gα subunit GPA1. However, co-expression of DRW2 with GPA1 produced luminescence, indicating DRW2 interacted with the Gα subunit GPA1 (Figure 5B). These data, in combination with our structural modeling studies, suggest that DRW2 is likely a non-canonical Gβ subunit.

**Figure 5.**
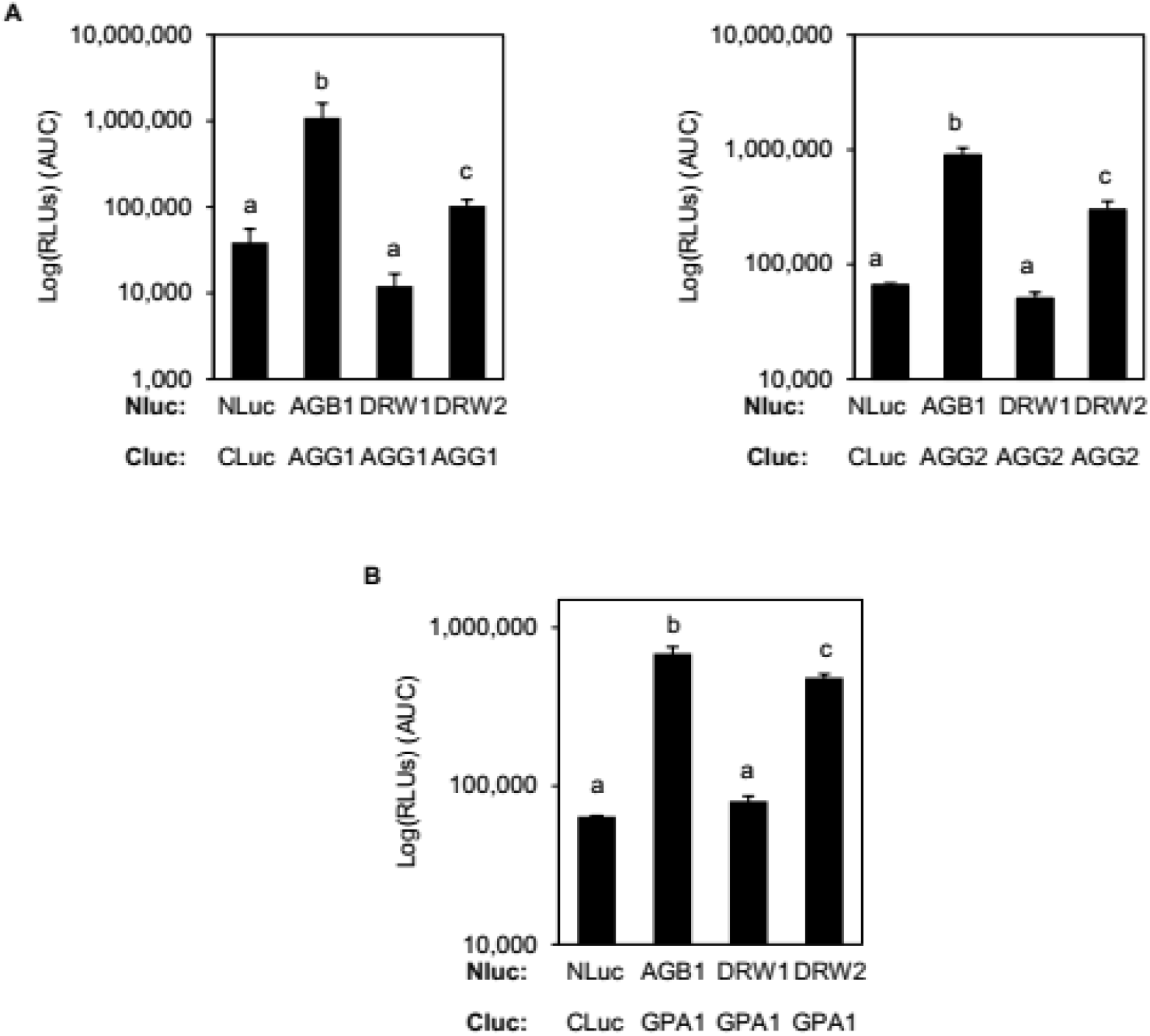
DRW2 interacts with Gγ subunits AGG1/AGG2 and the Gα subunit GPA1. (A) Quantitative assessment of protein-protein interactions between DRW1 and DRW2 with AGG1/2 using a reconstituted luciferase activity readout. NLuc-CLuc interaction is the negative control; AGB1-AGG1/2 and AGB1-AGG1/2 interactions are positive controls. Data represent median ± SD of four replicates. Different letters denote statistically significant differences (*P* < 0.05, two-tailed *t* test). (B) Quantitative assessment of protein-protein interactions between DRW1 and DRW2 with GPA1 using a reconstituted luciferase activity readout. NLuc-CLuc interaction is the negative control; AGB1-AGG1/2 and AGB1-AGG1/2 interactions are positive controls. Data represent median ± SD of four replicates. Different letters denote statistically significant differences (*P* < 0.05, two-tailed *t* test).

### Loss of DRW1 and DRW2 increases the levels of MAPK activation upon flagellin treatment

In order to understand the function of DRW1 and DRW2 in immunity, we wanted to investigate the phenotypes of the *drw1* and *drw2* loss-of-function mutants. To do this, we isolated two T-DNA insertion mutant alleles in *DRW1* and one T-DNA insertion mutant in *DRW2* that were obtained from the Arabidopsis Biological Resource Center (Figure S1A). The two T-DNA insertion alleles of *DRW1*, which we named *drw1-1* and *drw1-2*, have T-DNA insertions in the fourth and ninth exons, respectively (Figure S1A). The *drw2* mutant, *drw2-1*, has a T-DNA insertion in the 5’ untranslated region of *DRW2* (Figure S1A). To test if these mutants were loss-of-function mutations, we performed qRT-PCR to quantify *DRW1* and *DRW2* transcript levels in the *drw1-1* and *drw2-1* mutants (Figure S1B). Both the *drw1-1* and *drw2-1* single mutants showed very low levels of *DRW1* and *DRW2* transcripts in their respective mutants, suggesting that they are gene knockdown mutants (Figure S1B).

The heterotrimeric G proteins activate immune signaling downstream of the FLS2 receptor (Liu et al., 2013). To test if DRW1 or DRW2 participate in signaling downstream of FLS2, we performed MAPK activation assays on the *drw1* and *drw2* mutants. We measured MAPK activation 5 minutes post-elicitation with physiological levels of 100 nM flagellin. The *drw1-1, drw1-2*, and *drw2-1* single mutants showed an increase in MAPK activation upon flg22 treatment (Figure 6A). We performed semi-quantitation analysis on total activated MAPK levels in these mutants, and the *drw1* and *drw2* single mutants exhibited elevated levels of MAPK activation in response to flg22 treatment (Figure 6B). Our MAPK data suggests that DRW1 and DRW2 negatively affect MAPK activation in response to physiological concentrations of flagellin.

**Figure 6.**
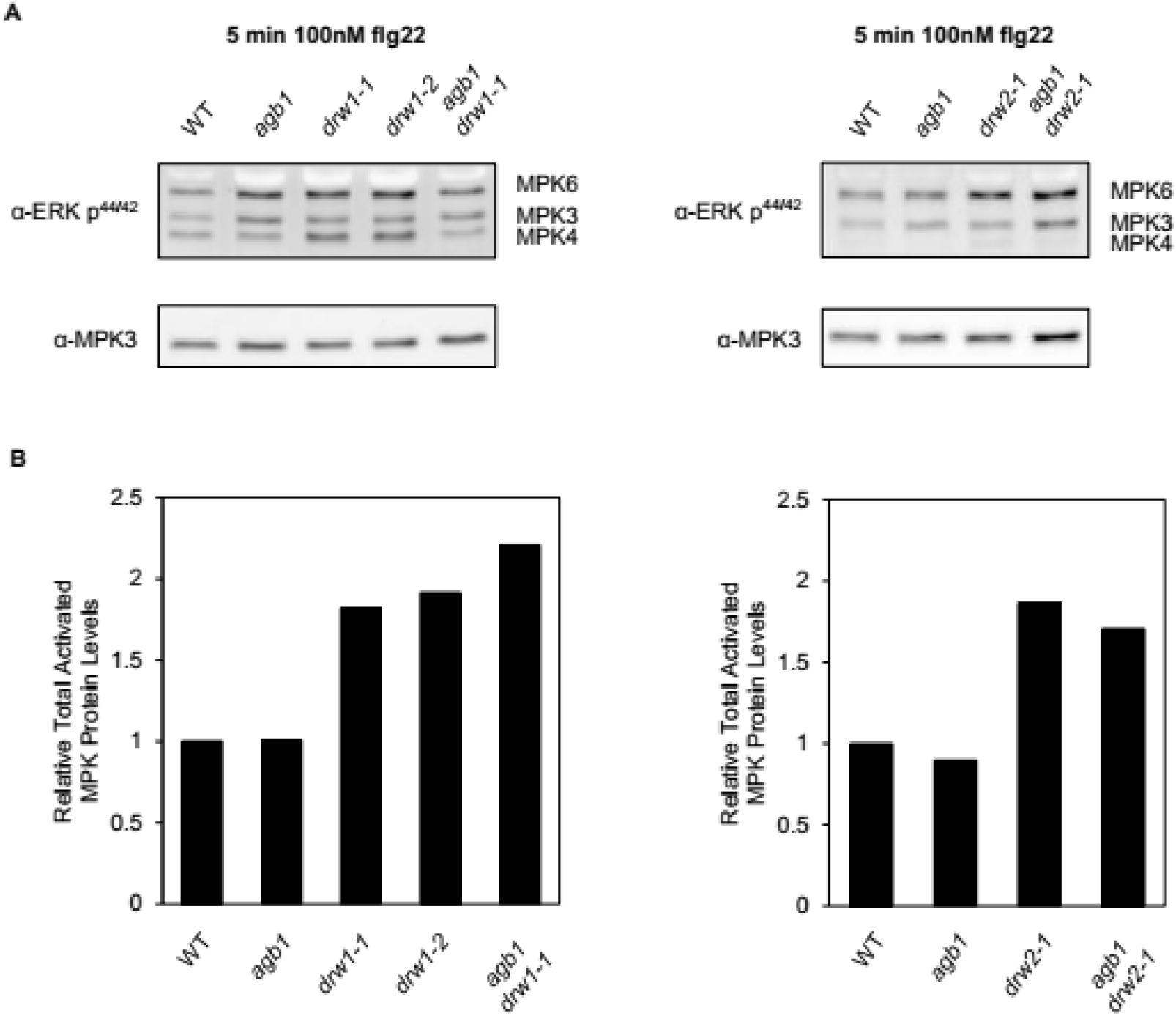
Loss of DRW1 or DRW2 increases MAPK activation upon flg22 elicitation. (A) Immunoblot analysis of phosphorylated, active MAPKs in 9-day-old WT*, drw1-1, drw1-2, agb1 drw1-1, drw2-1,* and *agb1 drw2-1* in response to 100 nM flg22 for 5 minutes. (B) Semi-quantitation of MAPK immunoblots in A. Data represent the mean of two replicates.

The genetic relationship between *DRW1*, *DRW2*, and *AGB1* is unknown. To better understand the genetic relationship between *AGB1*, *DRW1*, and *DRW2*, we created *agb1 drw1-1* and *agb1 drw2-1* double mutants and measured MAPK activation upon flagellin treatment. Interestingly, while AGB1 is known to be involved in MAPK activation upon flagellin treatment, we did not observe any changes in MAPK activation in the *agb1* mutant, as was observed previously (Liu et al., 2013). Furthermore, the *agb1 drw1-1* and *agb1 drw2-1* double mutants had similar elevated MAPK activation levels as the *drw1* and *drw2* single mutants in response to flagellin (Figure 6A and B). The lack of a phenotype in the *agb1* single mutant makes the determination of whether *AGB1*, *DRW1*, and *DRW2* function in the same or different pathways for MAPK activation difficult, and our double mutant analyses do not eliminate either possibility.

DRW1 and DRW2 share 89% sequence identity and similar elevated levels of MAPK activation in response to flagellin. It remains unclear if DRW1 and DRW2 function in the same pathway to repress the MAPK cascade in response to flagellin. To test this hypothesis, we created a *drw1-1 drw2-1* double mutant and measured the MAPK activation in response to flagellin treatment. The *drw1 drw2* double mutant had increased levels of MAPK activation compared to wild type and the *agb1* mutant (Figure 7A). We performed semi-quantitation on total activated MAPK levels in the *drw1*, *drw2*, and the *drw1 drw2* mutants. Activated MAPK levels in the *drw1* and *drw2* single mutants were similar to the *drw1 drw2* double mutant (Figure 7B). This data suggests that DRW1 and DRW2 may function in the same pathway, possibly in a non-redundant manner to repress MAPK activation upon flagellin treatment.

**Figure 7.**
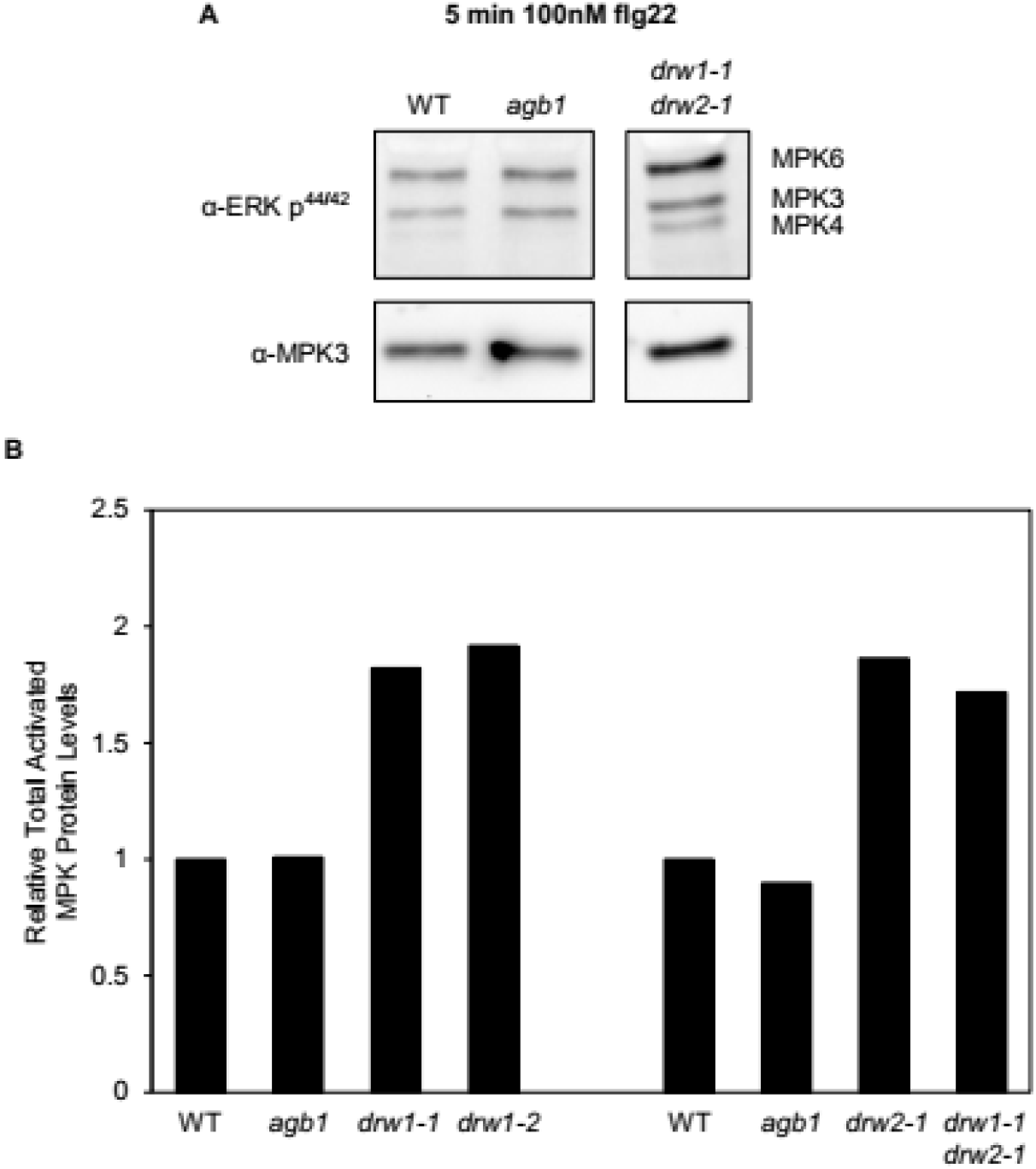
*drw1 drw2* double mutant increases levels of MAPK activation in response to flg22 treatment. (A) Immunoblot analysis of phosphorylated, active MAPKs in 9-day-old WT*, agb1,* and *drw1-1 drw2-1* in response to 100 nM flg22 for 5 minutes. (B) Semi-quantitation of MAPK immunoblots in A and Figure 6. Data represent the mean of two replicates.

### DRW1 and DRW2 are involved plant immunity as potential negative regulators of immunity

Our data suggests that DRW1 and DRW2 negatively regulate immunity upon PAMP treatment, in contrast to AGB1 which positively regulates immunity. However, it remains unclear whether DRW1 and DRW2 negatively regulate plant immunity upon pathogen infection. To test if DRW1 and DRW2 negatively regulate immune signaling, we infected plant leaves with the fungal pathogen *Alternaria brassicicola* and measured the lesion diameter. As expected, the lesion diameter in the *agb1* mutant was increased upon infection by *A. brassicicola* when compared to wild type (Figure 8A). The lesion diameters of the *drw1-1* and *drw2-1* single mutants were greater than the wild type lesion diameter but smaller than the lesion diameter of the *agb1* mutant. Interestingly, the *agb1 drw1-1* and *agb1 drw2-1* double mutants exhibited restored *A. brassicicola* susceptibility back to wild-type levels (Figure 8A). In contrast, the *drw1-1 drw2-1* double-mutant exhibited an increased resistance to *A. brassicicola* infection compared to wild type, which is in agreement with the MAPK activation experiments above (Figure 8A and 7). This suggests that DRW1 and DRW2 act in the same immune signaling pathway, possibly non-redundantly, in response to the fungal pathogen *A. brassicicola*. In light of this, the phenotypes of the *agb1 drw1-1* and *agb1 drw2-1* suggests that the DRWs and AGB1 may function in separate fungal immunity pathways. However, the increase in susceptibility in the *drw1-1* and *drw2-1* single mutants complicates this, and suggests that additional work is necessary to solve this discrepancy.

**Figure 8.**
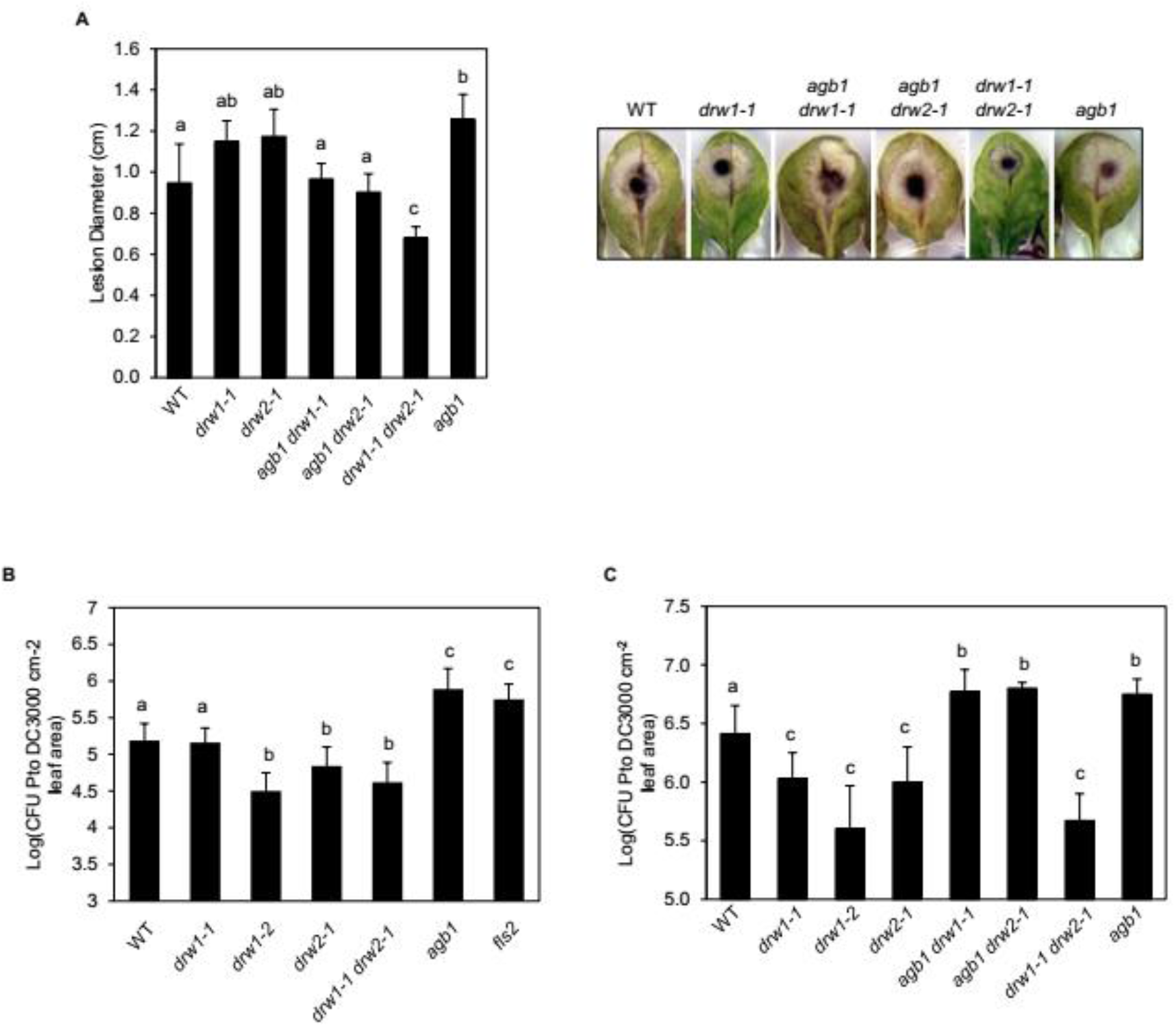
DRW1 and DRW2 negatively regulate plant immunity. (A) (Left) Quantitation of lesion development of 4-5-week-old leaves 3 days after inoculation with *A. brassicicola*. 4-5-week-old plant leaves were drop-inoculated with 5 µL of 5×10^5^ *Alternaria brassicicola* spores mL^-1^ for 3 days and lesion diameter was measured. Different letters in graph indicate significant differences (*P*-value <0.05, two-tailed *t* test). (Right) Representative pictures of leaves 3 days post-infection. (B-C) Growth analysis of bacterial pathogen *Pto* DC3000 in 4-5-week-old surface-inoculated leaves with *Pseudomonas syringae Pto* DC3000 (OD=0.0002) in the presence of 0.0075% Silwet and incubated the leaves on water-agar plates for 3 days. Data represent mean ± SD of six replicates. Different letters in (*B-C*) indicate significant differences (*P*-value <0.05, two-tailed *t* test).

Our data indicate that DRW1 and DRW2 negatively regulate immunity in response to fungal infection. However, it remains unknown whether DRW1 and DRW2 also negatively regulate immune signaling in response to bacterial infection. To determine if DRW1 and DRW2 negatively regulate immunity against bacterial pathogens, we infected plant leaves with the bacterial pathogen *Pseudomonas syringae Pto* DC3000 and then measured the number of bacteria growing in the extracted leaves. The *drw1-2* and *drw2-1* single mutants exhibited an increased resistance to *Pto* DC3000 compared to those in wild type and the susceptible *agb1* mutant (Figure 8B and C). The *agb1 drw1-1* and *agb1 drw2-1* double mutants were as susceptible as the *agb1* mutant (Figure 8C). The *drw1 drw2* double mutant was resistant to *Pto* DC3000 infection similar to the *drw1* and *drw2* single mutants (Figure 8B and C). This suggests that *DRW1* and *DRW2* are negative regulators of immunity that function in the same immune signaling pathway. Furthermore, our data suggests that *AGB1*, *DRW1*, and *DRW2* regulate bacterial immunity through the same pathway, and that *AGB1* is downstream of *DRW1* and *DRW2*.

## DISCUSSION

Plants encode canonical and non-canonical Gα and Gγ subunits. However, only one canonical Gβ subunit exists, severely limiting the number of heterotrimeric G protein complexes that can potentially function in different signal transduction pathways. The discovery of the non-canonical Gα and Gγ subunits opens the possibility that plants may also encode non-canonical Gβ subunits. In this study, we identify a WD40-repeat gene family named *DEFENSE REGULATED WD40-REPEAT* (*DRW*). This gene family consists of five genes (*DRW1-5*), of which *DRW1* and *DRW2* were transcriptionally repressed upon either bacterial or fungal pathogen infections. Protein structure homology analyses showed that DRW1 and DRW2 were predicted to form a β-propeller that is similar to AGB1. Moreover, DRW2 was predicted to form an N-terminal α-helix, serving as a coiled-coil domain. Since no protein structure is available of the plant Gβ subunit, future structural studies of AGB1, DRW1, and DRW2 interacting with a Gγ subunit are necessary to validate these predicted protein structures.

The heterotrimeric G protein complex localizes and functions at the plasma membrane (Oldham and Hamm, 2008; Temple and Jones, 2007). DRW2 co-localized to the plasma membrane with the Gγ subunit AGG1 and the pattern recognition receptor FLS2. Moreover, DRW2 interacted with the canonical Gα and Gγ subunits GPA1 and AGG1/AGG2, respectively. Co-immunoprecipitation experiments are needed to validate the protein-protein interactions of DRW2 with the canonical heterotrimeric G proteins that we observed in the protoplast experiments. Small populations of DRW1 localized to the plasma membrane, but DRW1 did not interact with GPA1 or AGG1 and AGG2. However, DRW1 may interact with the non-canonical Gα (XLG1/2/3) and Gγ (AGG3) subunits rather than the canonical G proteins, although further studies are needed.

In accordance with the transcriptional expression data, gene knockdown mutants of *DRW1* or *DRW2* had increased MAPK activation in response to flagellin treatment, unlike the *agb1* mutant which was similar to wild type. Moreover, the *drw1* and *drw2* mutants had increased resistance to *P. syringae Pto* DC3000. When both *DRW1* and *DRW2* are knocked down, these plants exhibit a broad-spectrum resistance to both fungal and bacterial pathogen infections. Our data show that DRW1 and DRW2 function to negatively regulate immune signaling.

Plant immunity is tightly regulated and fine-tuned to prevent autoimmunity (Couto and Zipfel, 2016). Plants employ many negative regulators to inactivate immune signaling. Our data indicate that DRW1 and DRW2 are negative regulators of immunity. However, the mechanism by which DRW1 and DRW2 regulate immunity remains a mystery. Further research is necessary to elucidate the mechanism by which DRW1 and DRW2 function in immunity. For example, gene knockdown of *DRW1* and *DRW2* caused an increase in resistance to pathogen infection, but it would be interesting to see if transgenic plants overexpressing *DRW1* and/or *DRW2* in the wild-type background would exhibit increased susceptibility upon pathogen infection. This would validate our hypothesis that DRW1 and DRW2 are negative regulators of immunity. Moreover, the genetic relationship between *DRW1*, *DRW2*, and *AGB1* is complex as the *agb1 drw1* and *agb1 drw2* double mutants suppress the *agb1* susceptibility phenotype to the fungal pathogen *A. brassicicola* but not against the bacterial pathogen *P. syringae*. Illuminating the genetic relationship between *DRW1*, *DRW2*, and *AGB1* may be the key to determine the mechanism by which DRW1 and DRW2 function in immunity. In order to better understand this relationship, an *agb1 drw1 drw2* triple mutant would need to be created to identify other immunity phenotypes not seen in the double mutants.

Plants encode over 200 WD40-repeats which provides a challenge in identifying non-canonical Gβ subunits (van Nocker and Ludwig, 2003). A reverse genetic screen to identify non-canonical Gβ subunits would be massive and labor-intensive. However, our study provides a framework that can be employed to identify potential Gβ subunits in other organisms. Transcriptional analysis paired with homology modeling narrows the large list of WD40-repeat proteins to a feasible number for characterization. However, this method requires a sequenced genome for the organism of interest. Furthermore, transcriptional analyses under different abiotic and biotic conditions are also required for this method. As additional crop genomes are released and biotic stress transcriptional profiles are generated, this methodology to identify Gβ subunits will uncover additional non-canonical Gβ subunits in agricultural species that will allow us to minimize crop loss due to pathogens.

## MATERIALS AND METHODS

### Identification of seven WD repeat orthologues

*Arabidopsis thaliana* WD40 protein annotations (N=358) were downloaded from the WD40-repeat protein Structure Predictor database (http://wu.scbb.pkusz.edu.cn/wdsp/index.jsp; Wang et al., 2015). From these annotations, 262 proteins were identified as having exactly seven WD repeats and a subset of 168 proteins with known expression profiles were used for further analysis. Protein sequences were aligned in MUSCLE and maximum likelihood trees were generated based on the JTT matrix-based model in MEGA7 (Kumar et al., 2016). Initial tree(s) for the heuristic search were obtained automatically by applying Neighbor-Join and BioNJ algorithms to a matrix of pairwise distances estimated using a JTT model, and then selecting the topology with superior log likelihood value. All amino acid positions with less than 90% site coverage were eliminated, leaving a total of 266 positions in the final dataset.

### Plant Materials and Growth Conditions

Surface-sterilized seeds of *Arabidopsis thaliana* accession Columbia-0 (Col-0) were stratified for at least 2 days and sown in 12-well microtiter plates sealed with parafilm. Each 12-well plate contained 12 seedlings with 1 of filter-sterilized 0.5X MS liquid (pH 5.7–5.8) [4.43 g/L Murashige and Skoog basal medium with vitamins (Murashige and Skoog, 1962) (Phytotechnology Laboratories, Shawnee Missions, KS), 0.05% (w/v) MES hydrate, 0.5% (w/v) sucrose], respectively. Alternatively, surface-sterilized and stratified seeds were sown on MS agar plates [0.5X MS, 0.75% (w/v) agar (PlantMedia, Chiang Mai, Thailand)] and sealed with parafilm. Unless otherwise stated, plates were placed on grid-like shelves over water trays on a Floralight cart (Toronto, Canada), and plants were grown at 21°C and 60% humidity under a 12 hr light cycle (70–80 μE m^-2^ s^-1^ light intensity). Unless otherwise stated, media in microtiter plates were exchanged for fresh media on day 7. For bacterial infection experiments, *Arabidopsis* plants were grown on soil [3:1 mix of Fafard Growing Mix 2 (Sun Gro Horticulture, Vancouver, Canada) to D3 fine vermiculite (Scotts, Marysville, OH)] at 22°C daytime/18°C nighttime with 60% humidity under a 12 hr light cycle (100 µE m^-2^ s^-1^ light intensity). *Nicotiana benthamiana* plants were grown on soil (3:1 mix) on a Floralight cart at 22°C under a 12 hr light cycle (100 µE m^-2^ s^-1^ light intensity) for 4 weeks.

The following homozygous Col-0 T-DNA insertion lines and mutants were obtained from the Arabidopsis Biological Resource Center (ABRC, Columbus, Ohio): *agb1* (CS3976)*, drw1-1* (SALK_142665C), *drw1-2* (SALK_098040C), *drw2-1* (WiscDsLox3E04/CS849082), *fls2* (SAIL_691_C4).

### Vector Construction and Transformation

To generate estradiol-inducible C-terminally tagged *GFP* and *RFP* (*XVE:X-GFP/RFP*) DNA constructs, *attB* sites were added via PCR-mediated ligation to the coding sequences of cDNAs, and the modified cDNAs were recombined into pDONR221 entry vector and then into pABindGFP and pABindRFP destination vectors (Bleckmann et al., 2010), according to manufacturer’s instructions (Gateway manual; Invitrogen, Carlsbad, CA). Transient expression of *XVE:X-GFP/RFP* constructs in *N. benthamiana* leaves was performed as previously described (Bleckman et al., 2010) with the following modification: transformed *Agrobacterium* strains were grown in LB medium supplemented with 50 µg/mL rifampicin, 30 µg/mL gentamycin, 50 μg/mL kanamycin and 100 µg/mL spectinomycin, in the absence of a silencing suppressor, to an OD_600_ of 0.7. Transgene expression was induced 4-8 hr (for microscopy) after spraying with 20 µM β-estradiol and 0.1% Tween-20.

For split-luciferase assays, *GPA1*, *AGG1*, and *AGG2* were inserted in the pUC19-35S∷CLuc vector from (Chen et al., 2008). *AGB1*, *DRW1*, and *DRW2* were inserted in the pUC19-35S∷NLuc vector from (Chen et al., 2008).

### Confocal Microscopy

4-week-old *N. benthamiana* leaves were imaged using a 40X 1.0 numerical aperture Zeiss water-immersion objective and a Zeiss LSM 510 Meta confocal microscopy system. GFP and RFP were excited with a 488-nm argon laser and 561-nm laser diode, respectively. GFP and RFP emissions were detected using a 500-550 nm and 575-630 nm filter sets, respectively. Plasmolysis was induced by 5-10 min treatment of *N. benthamiana* leaf strips with 0.8 M mannitol, and co-localization of GFP/RFP-tagged proteins to Hechtian strands was made visible by over-exposing confocal images using ZEN software.

### MAPK Activation Assay

9-day-old seedlings were elicited with 100 nM flg22 for 5, 15, and/or 30 min. MAPK activation assay was performed as previously described (Lawerence et al., 2017). 20 μl of supernatant was loaded onto a 10% SDS-PAGE gel, and the separated proteins were transferred to PVDF membrane (Millipore) and probed with phosphor-p44/p42 MAPK (Cell Signaling Technology, Danvers, MA) and MPK3 antibodies (Sigma-Aldrich, St. Louis, MO) at 1:2000 dilution in 5% (w/v) nonfat milk in 1X PBS. The combined signal intensities of phosphorylated MPK3/4/6 were quantified using NIH ImageJ and normalized to that of total MPK3 (loading control).

### Split-Luciferase Complementation Assay

Arabidopsis protoplasts were isolated and transfected as previously described (Sheen (http://genetics.mgh.harvard.edu/sheenweb/). In brief, 3-4 week-old Arabidopsis plants were cut into 0.5-1.0 mm leaf strips and incubated in enzyme solution (400 mM mannitol, 20 mM KCl, 20 mM MES pH 5.7, 100 mg Cellulase R10, 20 mg Macerozyme, 10 mM CaCl_2_, 0.1% BSA [Sigma A7906] sterile filtered) at room temperature for 4-8 hr. 1X volume of cold W5 solution (30.8 mM NaCl, 125 mM CaCl_2_, 5 mM KCl, 2 mM MES pH 5.7, 50 mM glucose) was added to the enzyme solution and filtered through a 20 μm nylon mesh into a polystyrene test tube. Samples were centrifuged at 100 x g for 2 minutes, the supernatant was removed, then cells were washed twice with cold W5 solution and suspended in 3 mL cold W5 solution. Protoplasts were quantified using a hemocytometer and resuspended in cold MMg solution (400 mM mannitol, 15 mM MgCl_2_, 4 mM MES pH 5.7) to yield a concentration of 5×10^5^cells mL^-1^. 10 μg of each vector was mixed with 200 μL of 5 x 10^5^ cells mL^-1^ and gently mixed. 1X volume of PEG solution was added (200 mM mannitol, 100 mM CaCl_2_, 40% (w/v) PEG 4000) to samples, tubes were gently inverted 10 times, and then incubated at room temperature for 15 minutes. 1 mL of W5 solution was added to each sample and gently mixed. Samples were then centrifuged at 100 x g for 2 minutes, the supernatant was removed and the cells were washed with 1 mL W5 solution. Samples were centrifuged again at 100 x g for 1 minute and 900 μL of supernatant was removed. Protoplasts were transferred to 6-well plates, coated with 10% calf-bovine serum, with 1 mL W5 solution. Protoplasts were incubated under constant light for 18-24 hr.

Transfection efficiency was determined by the number of protoplasts expressing citrine using a Zeiss (Oberkochen, Germany) AxioObserver D1 fluorescence microscope under UV illumination with Filter Set 52 (excitation filter 488/20 nm; dichroic mirror 505 nm; emission filter 530/50 nm). Protoplasts were centrifuged at 1000 rpm for 2 minutes, approximately 900μL of supernatant removed, and transferred to luminometer cuvettes. 100μL of luciferin (Reconstituted Promega Luciferin Assay Buffer) was added to each cuvette and immediately measured using a luminometer (Berthold FB12 Single Tube Luminometer) for 6 minutes. The area under the curve was calculated.

### Bacterial Pathogen Infection Assay

Pathogen assays on 4- to 5-week-old adult leaves were performed as previously described (Chezem et al., 2017). In brief, *Pseudomonas syringae* pv*. tomato* DC3000 (*Pto* DC3000) was grown overnight in LB and 25 µg/mL rifampicin (Sigma-Aldrich) and then washed in sterile water twice. *P. syringae* was resuspended in water to the desired OD_600_ and adult leaves of 4- to 5-week-old plants were surface-inoculated with the bacterial inoculum (OD_600_ = 0.002 or 10^6^ colony-forming units (CFU)/cm^2^ leaf area) in the presence of 0.0075% Silwet L-77 (Phytotechnology Laboratories) and incubated on 0.8% (w/v) tissue-culture water agar plates for 4 days. Leaves were surface-sterilized in 70% ethanol, washed in sterile water, and dried on paper towels. Bacteria were extracted into water, using an 8-mm stainless steel bead and a ball mill (25 Hz for 3 min). Serial dilutions of the extracted bacteria were plated on LB agar plates for CFU counting.

### Fungal Pathogen Infection Assay

*Alternaria brassicicola* strain FSU218 (Fungal Reference Center, Jena, Germany) was used for fungal infections. *A. brassicicola* was grown on PDA (1% Potato Dextrose Agar) plates at 21°C, 16 hr photoperiod, <100 µE m^-2^ s^-1^, wrapped in parafilm to maintain high humidity for 3 weeks before collecting spores. *A. brassicicola* conidia spores were harvested and resuspended in sterile water, and incubated at RT for 24 hr. Conidia were quantified using a hemocytometer and the spore inoculum was adjusted to a concentration of 5×10^5^ spores mL^-1^. 5 μL droplets were placed on the surface of detached leaves and leaves were incubated at 21°C, 16-hr photoperiod, >100 μE m^-2^ s^-1^, in high humidity for 3 days before imaging leaves.

### RNA isolation and quantitative PCR (qPCR)

Total RNA was extracted into 1 mL of TRIzol reagent (Invitrogen) according to manufacturer’s instructions. 2 µg of total RNA was reverse-transcribed with 3.75 µM random hexamers (Qiagen, Hilden, Germany) and 20 U of ProtoScript II (New England Biolabs, Boston, MA). The resulting cDNA:RNA hybrids were treated with 10 U of DNase I (Roche) for 30 min at 37°C, and purified on PCR clean-up columns (Macherey-Nagel, Düren, Germany). qPCR was performed with Kapa SYBR Fast qPCR master mix (Kapa Biosystems, Wilmington, MA) and CFX96 or CFX384 real-time PCR machine (Bio-Rad, Hercules, CA). The thermal cycling program is as follows: 95°C for 3 min; 45 cycles of 95°C for 15 sec and 53°C or 55°C for 30 sec; a cycle of 95°C for 1 min, 53°C for 1 min, and 70°C for 10 sec; and 50 cycles of 0.5°C increments for 10 sec. Biological replicates of control and experimental samples, and three technical replicates per biological replicate were performed on the same 96- or 384-well PCR plate. Averages of the three Ct values per biological replicate were converted to differences in Ct values relative to that of control sample. Pfaffl method (Pfaffl 2001) and calculated primer efficiencies were used to determine the relative fold increase of the target gene transcript over the housekeeping *eIF4AI* gene transcript for each biological replicate.

## ACKNOWLEDGEMENTS

We thank Josh Gendron for help with interpreting the data. This work was supported by T32 GM007223 (to J.C.M).

---

**Supplementary Figure 1.**
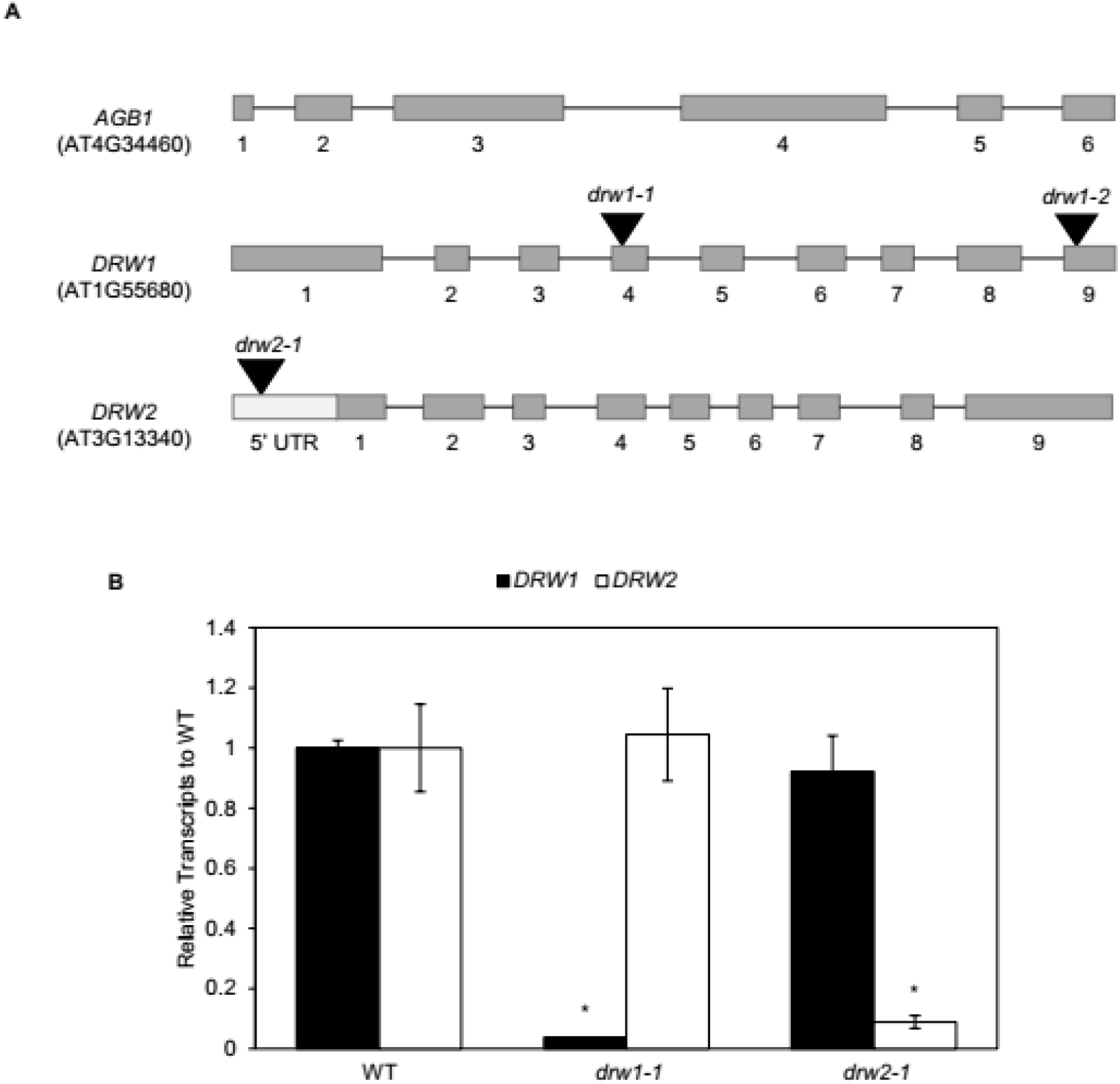
*DRW1* and *DRW2* gene models and qPCR. (A) Gene models of DRW1 and DRW2 marked with the approximate locations of the T-DNA insertions. Grey boxes indicate exons and the black lines indicate introns. Light grey box represents the 5’ untranslated region (UTR). (B) qPCR analysis of *DRW1* and *DRW2* transcripts in wild-type and their respective backgrounds.

